# Myocyte Enhancer Factor 2A Orchestrates Vascular Redox Homeostasis via Direct Transcriptional Activation of SIRT1

**DOI:** 10.1101/2025.07.15.665025

**Authors:** Benrong Liu, Lei Fang, Chunxia Miao, Xinyu Wen, Xiumiao Zheng, Minxing Xu, Junli Lin, Yujuan Xiong, Shi-Ming Liu

## Abstract

Myocyte enhancer factor 2A (MEF2A), a transcription factor implicated in coronary artery disease, remains unexplored in vascular redox regulation. To address this gap and overcome limitations of current antioxidant therapies, we investigated MEF2A’s role in oxidative defense using human umbilical vein endothelial cells (HUVECs) and murine models. Adenoviral vectors encoding MEF2A-specific shRNA or mRNA were employed to silence or overexpress MEF2A in HUVECs. For in vivo validation, endothelial-targeted MEF2A knockdown was achieved via AAV1-shRNA delivery in mice fed a high-fat diet. Systemic redox status was assessed by measuring reactive oxygen species (ROS), glutathione homeostasis (GSH/GSSG ratio), NADH/NAD^+^ balance, mitochondrial membrane potential (ΔΨm), and 8-hydroxy-2′-deoxyguanosine (8-OHdG). Mechanistic insights were derived from immunofluorescence, qPCR, western blotting, and dual-luciferase reporter assays. MEF2A silencing induced redox imbalance, characterized by elevated ROS, reduced GSH/GSSG ratio, and ΔΨm collapse. Conversely, MEF2A overexpression synergized with SIRT1 to restore glutathione pools, maintain NAD^+^ homeostasis, and suppress ROS under oxidative stress. Chromatin immunoprecipitation confirmed direct MEF2A binding to two cis-elements in the SIRT1 promoter, driving transcriptional activation. In vivo, MEF2A-deficient mice exhibited amplified vascular oxidative damage, including elevated DNA damage marker (8-OHdG) and ROS levels. Downregulation of SIRT1/PGC-1α in MEF2A silenced cells was verified in vivo. Our findings establish MEF2A as a master regulator of endothelial redox defense via the SIRT1-PGC-1α axis, offering a mechanistic foundation for targeting oxidative cardiovascular disorders. This work suggests pharmacological MEF2A activation as a novel strategy for precision antioxidant therapy in vascular medicine.

## Introduction

Oxidative stress—a pathophysiological state characterized by excessive reactive oxygen species (ROS) production overwhelming endogenous antioxidant defenses—serves as a critical driver of cellular dysfunction and tissue damage across chronic diseases[1]. In the cardiovascular system, this redox imbalance triggers endothelial injury, initiating a cascade of events that culminate in atherosclerosis, hypertension, and ischemic heart disease[2–5]. While ROS are essential signaling molecules at physiological levels, their pathological accumulation induces DNA damage, lipid peroxidation, and mitochondrial impairment, with vascular endothelial cells being particularly vulnerable due to their direct exposure to hemodynamic stressors[6,7].

Vascular endothelial cells, far beyond their traditional role as passive blood vessel linings, function as dynamic biosensors regulating vascular tone, thromboresistance, and inflammatory responses through paracrine signaling[8]. Emerging evidence positions endothelial redox dysregulation as the “first domino” in atherosclerosis development, where oxidative inactivation of nitric oxide synergizes with oxidized LDL uptake to drive foam cell formation and plaque progression[8–10]. Despite advances in antioxidant therapies (e.g., vitamins, Probucol and Related Phenols), clinical translation remains hampered by systemic toxicity and inability to target endothelial-specific pathways[11]. This therapeutic gap underscores the urgency to identify master transcriptional regulators of endothelial redox homeostasis. Sirtuin 1 (SIRT1), an NAD^+^-dependent deacetylase, and peroxisome proliferator-activated receptor γ coactivator-1α (PGC-1α), a master regulator of mitochondrial biogenesis, form a critical nexus for redox defense [12]. SIRT1 deacetylates PGC-1α to enhance mitochondrial function and ROS scavenging [13]. Pharmacological activators like resveratrol improve vascular function[14], yet their reliance on upstream transcriptional control mechanisms remains unexplored. This gap parallels broader failures of nonspecific antioxidants, highlighting the urgency to define transcription factors coordinating mitochondrial quality control and endogenous defenses.

Myocyte enhancer factor 2A (MEF2A), a member of the MADS-box transcription factor family, has emerged as a paradoxical player in cardiovascular biology [15–17]. Originally identified for its essential role in muscle development through regulating myocyte differentiation and morphogenesis [18], MEF2A has since been shown to exhibit pleiotropic functions across tissues. In immune regulation, a study by Renato Ostuni’s group revealed that PGE2 signaling in macrophages functionally inactivates MEF2A to modulate inflammatory gene expression, demonstrating its context-dependent role in immune modulation[19]. In developmental biology, MEF2A regulates cell differentiation in Dictyostelium discoideum, where mef2A-deficient strains exhibit impaired prespore cell formation and spore production, highlighting its evolutionary conservation in cell-fate determination[20]. Emerging evidence also links MEF2A to metabolic regulation, with studies showing its role in promoting cardiomyocyte proliferation through SphK2/S1P signaling, thereby extending the cardiac regenerative window [21]. Conversely, in cancer biology, MEF2A is overexpressed in colorectal cancer, where it activates WNT/β-catenin signaling to drive tumor proliferation, while metformin inhibits its oncogenic activity by promoting ubiquitin-mediated degradation[15]. Neurodegenerative disease research further reveals MEF2A’s involvement in autophagy regulation[22]; its downregulation via enhancer methylation correlates with impaired autophagic flux in Alzheimer’s disease models[23]. These multifaceted roles collectively position MEF2A as a central node integrating developmental, metabolic, and stress-responsive signaling pathways.

While human genetic studies link MEF2A mutations to premature coronary artery disease [24,25] and dilated cardiomyopathy[26], its mechanistic role in vascular redox regulation remains enigmatic. Our prior work revealed MEF2A’s capacity to delay H₂O₂-induced endothelial senescence via PI3K/Akt/SIRT1 signaling[27] and mediate resveratrol’s vasculoprotective effects[22]. These findings hint at MEF2A’s potential as a redox-sensitive transcription factor, yet key questions persist: Does MEF2A directly orchestrate antioxidant gene networks? What molecular effectors execute its protective functions?

Here, we bridge this knowledge gap by delineating a novel MEF2A-SIRT1-PGC-1α axis essential for endothelial redox homeostasis. Through integrated in vitro and in vivo approaches, we demonstrate that: (1) MEF2A deficiency precipitates mitochondrial ROS overflow and glutathione depletion via SIRT1 suppression; (2) MEF2A directly transactivates SIRT1 through conserved promoter elements, establishing a self-reinforcing antioxidant loop; (3) Endothelial-specific MEF2A knockdown in mice recapitulates human atherosclerotic phenotypes, including oxidative DNA damage and PGC-1α downregulation. Our findings not only redefine MEF2A as a transcriptional guardian of vascular redox equilibrium but also provide a mechanistic blueprint for developing precision therapies targeting this axis in cardiovascular diseases.

## Materials and Methods

### Cell Culture

Primary human umbilical vein endothelial cells (HUVECs) were isolated from human umbilical cord veins using a combination of trypsin-collagenase digestion and the differential adhesion method, followed by selection using endothelial cell-specific culture medium. HUVECs were maintained in EBM-2 basal medium (CC3156, Lonza, Basel, Switzerland) supplemented with 3% fetal bovine serum (FBS; Lonza, Basel, Switzerland), endothelial cell growth supplement (CC4176, Lonza, Basel, Switzerland), and antibiotics, and incubated at 37°C in a humidified 5% CO₂ atmosphere. The isolation protocol was approved by the Ethics Committee of the Second Affiliated Hospital of Guangzhou Medical University (Approval No.: 2021-LCYJ-ZF-06).

### Oxidative stress cell model

HUVECs were seeded in 48-well or 96-well plates and allowed to adhere overnight. The culture medium was then replaced with fresh complete medium containing a gradient of hydrogen peroxide (H₂O₂; 0, 50, 100, 200, or 400 μM). Cells were exposed to H₂O₂ for 1, 2, or 4 hours, after which the treatment medium was aspirated and replaced with fresh medium. Following an additional 24-hour incubation under standard culture conditions, cell viability was quantified. The exposure parameters yielding approximately 60% viability (200 μM H₂O₂ for 2 hours) were selected to establish the oxidative stress model for subsequent experiments.

### Silencing or Overexpression of MEF2A in Vitro

Adenoviral expression vectors containing MEF2A/SIRT1-specific short hairpin RNA (shRNA) or overexpression cassettes were constructed. The mature siRNA sequence of MEF2A and SIRT1 shRNA are 5’-GGGCAGUUAUCUCAGGGU U-3’ and 5’- GGCGGCUUGAUGGUAAUCAGUA-3’, respectively. A green fluorescent protein (GFP) reporter was incorporated downstream of target genes for expression monitoring. Viral packaging, purification, and titer determination were performed by Shandong Weizhen Biosciences Co., Ltd. Control groups received mir30-shRNA or GFP-expressing adenoviruses. Transduction efficiency was verified by fluorescence microscopy prior to functional assays.

### Assessment of GSH/GSSG Ratio

The glutathione redox status was determined using a commercial GSH/GSSG Assay Kit (S0053; Beyotime, Shanghai, China) according to manufacturer protocols. Samples from cells, plasma, and vascular tissues were processed in parallel.

### Detection of Cellular and Mitochondrial Reactive Oxygen Species (ROS)

Intracellular ROS was quantified using dihydroethidium (DHE; S0063, Beyotime, Shanghai, China). Cells were washed with PBS (37°C), incubated with 5 μM DHE in serum-free medium (30 min, dark), then counterstained with Hoechst 33342 (10 min). Fluorescence intensity (Ex/Em: 535/610 nm) was quantified using Image J.

Mitochondrial ROS was assessed using BBcellProbe® OM08 (BB-46091; Bestbio, Nanjing, China). Cells were incubated with 1:100-diluted probe in HBSS (30 min, dark), washed, and imaged immediately. Quantification used identical parameters across experimental groups.

### NADH/NAD^+^ Ratio Detection

The NAD^+^ and NADH contents in cells were detected using the NADH/NAD^+^ Detection Kit (WST - 8 method) (S0175) from Beyotime, and then the NADH/NAD^+^ ratio was calculated. The detection steps were carried out according to the kit instructions and are briefly described as follows: For the prepared cell samples, after being washed with PBS, cell lysis buffer was added to lyse the cells. Next, a heat - treated sample for NADH determination and a sample directly used for the determination of the total amount of NAD^+^ and NADH were prepared respectively. Then, ethanol dehydrogenase working solution was added to both samples and incubated in the dark. Finally, after adding the chromogenic solution and incubation, the OD value was measured at 450 nm, and the total amount of NAD^+^ and NADH and the content of NADH were calculated according to the standard curve. The content of NAD^+^ was obtained by subtracting the separately measured NADH amount from the total amount of NAD^+^ and NADH.

### Mitochondrial Membrane Potential Assay

Tetramethylrhodamine Ethyl Ester (TMRE; C2001S, Beyotime, Shanghai, China) staining was performed as previously described[22], in brief: Cells were incubated with TMRE working solution (15-45 min, dark), washed with HBSS, then counterstained with Hoechst 33342 (5 min). Fluorescence images were captured within 30 min post-staining.

### Dual-Luciferase Reporter Assay

The SIRT1 promoter (-2000 to +66 bp) was cloned into pGL3-Basic (Promega, Madison, USA). Mutant constructs with disrupted MEF2A binding sites (predicted by LASAGNA-Search 2.0[28]) were generated by overlap PCR. Co-transfection into 293T cells with Renilla luciferase vector (phRL-TK; Promega, Madison, USA) was performed using Lipofectamine 3000. Luminescence was measured 48-h post-transfection using the Dual-Luciferase Reporter Gene Assay Kit (E1910, Promega, Madison, USA) with a GloMax® Explorer luminometer (Promega, Madison, USA).

### Chromatin Immunoprecipitation (ChIP)

ChIP analysis was performed using the SimpleChIP® Enzymatic Chromatin IP Kit (#9003; CST, Danvers, USA). Briefly, HUVECs were cross-linked with 1% formaldehyde (10 min, RT) followed by quenching with 0.125 M glycine (5 min, RT). Cells were harvested, washed twice with ice-cold PBS, and lysed in 1× ChIP lysis buffer supplemented with 1 mM DTT and 1× protease inhibitor cocktail (PIC). Chromatin was digested with micrococcal nuclease (MNase) at 37°C for 20 min to generate 150-300 bp fragments.

The digested chromatin was immunoprecipitated overnight at 4°C with 2 μg of ChIP-grade anti-MEF2A antibody (#9736; CST, Danvers, USA) or IgG control (#2729; CST, Danvers, USA) coupled to Protein A/G magnetic beads. After sequential washing with low-salt and high-salt buffers, bound complexes were eluted in ChIP elution buffer. Crosslinks were reversed by incubation with 200 mM NaCl at 65°C for 4 h, followed by proteinase K treatment (45 min, 45°C). DNA was purified using silica-membrane columns and quantified by qPCR with SYBR Green Master Mix (LightCycler^®^ 480, Roche, Rotkreuz, Switzerland). Primer sequences spanning predicted MEF2A binding sites are listed in Supplementary Tables (Table S1).

### Experimental Animals

Male C57BL/6J (6 weeks old; Guangdong Experimental Animal Center, Foshan, China) were housed under specific pathogen-free conditions (23±2°C, 60% humidity, 12-h light cycle). After AAV1 (1×10¹² Vg/mouse) delivery via tail vein, mice received high-fat diet (HF60; Research Diets) for 16 weeks. All procedures followed ARRIVE guidelines and were approved by the Institutional Animal Care and Use Committee of Guangzhou Medical University (No. 2021-LCYJ-ZF-06).

### Vascular-Specific Mef2a Knockdown

The endothelial-specific pAV[Icam2]-shMef2a-GFP AAV1 vector was engineered by packaging Icam2 promoter-driven Mef2a shRNA into AAV serotype 1 particles. Mef2a knockdown group (si-Mef2a, n = 20) received pAV[Icam2]-shMef2a-GFP AAV1 injection, and the control groups (si-NC, n = 20) received pAV[Icam2]-miR30-shRNA-GFP AAV1 containing scrambled sequences. Mice were administered 5 × 10^11^ viral genome (vg) particles via tail vein injection. Following viral transduction, animals were fed a high-fat diet (HF60; Research Diets) containing 60% fat for 16 weeks to induce atherosclerosis.

### Tissue Harvesting

Mice were anesthetized via inhalation of 2% isoflurane, with anesthesia depth confirmed by absence of toe pinch reflex. Following euthanasia through cervical dislocation, terminal blood collection was performed by orbital enucleation using heparinized capillaries. Major organs including aortic arch, carotid bifurcation, hepatic lobe, and cardiac ventricles were immediately dissected. All tissues were snap-frozen in liquid nitrogen or fixed in 4% paraformaldehyde (G1101, PFA, Servicebio, Wuhan, China) for downstream analysis

### Vascular ROS Detection

Fresh carotid artery cryosections (10 μm) were stained with 5 μM dihydroethidium (DHE; S0063, Beyotime, Shanghai, China) in PBS at 37°C for 30 min. After counterstaining with DAPI (1 μg/mL, 5 min) and PBS washes, sections were mounted with ProLong® Gold Antifade Reagent (P36930, Thermo Fisher Scientific, Waltham, USA). Fluorescence imaging was performed using a Zeiss LSM 880 confocal microscope (20× objective) with identical exposure settings across groups.

### Immunofluorescence Staining

Paraformaldehyde-fixed vessels were embedded in paraffin and sectioned at 5 μm thickness. Antigen retrieval was conducted in EDTA buffer (pH 8.0, 95°C, 20 min) followed by endogenous peroxidase blockade with 3% H₂O₂. Sections were blocked with 5% BSA for 1 h before incubation with anti-8-OHdG antibody (1:200; sc-393871, Santa Cruz Biotechnology, Dallas, USA) overnight at 4°C. Tyramide-FITC signal amplification (1:100) was performed after secondary antibody incubation. Nuclei were stained with DAPI (1 μg/mL, 5 min) prior to imaging on a Zeiss LSM 880 system.

### RNA Isolation and qPCR

Total RNA was isolated using TRIzol® (Invitrogen, Waltham, USA) with the following modifications for vascular tissues: Aortic segments were snap-frozen in liquid nitrogen and homogenized in a cryomill. Lysates were phase-separated by chloroform addition (1:5 v/v) and centrifuged at 12,000×g (4°C, 15 min). RNA precipitation used ice-cold isopropanol (1:1 v/v), followed by 75% ethanol washes. RNA integrity was verified by 1% agarose gel electrophoresis. Reverse transcription employed FastKing RT SuperMix (KR123; TIANGEN, Beijing, China) with genomic DNA removal. SYBR Green-based qPCR was performed on a LightCycler® 480 system (Roche, Rotkreuz, Switzerland). Primer sequences for human/mouse targets are listed in Supplementary Table 1.

### Western Blotting

Aortic proteins were extracted using RIPA buffer (50 mM Tris-HCl, pH 7.4; 1% NP-40; 0.5% sodium deoxycholate) supplemented with protease/phosphatase inhibitors (PIC, FD1002, Hangzhou Fdbio Technology, Hangzhou, China). Tissue homogenization used a Tissue Grinder at 60 Hz for 5 cycles (60 sec/cycle). Lysates were cleared by centrifugation (12,000×g, 20 min, 4°C).

Protein concentration was determined by BCA assay (23227; Thermo Scientific, Waltham, USA). Samples (30 μg/lane) were resolved on 10% SDS-PAGE gels and transferred to PVDF membranes (IPFL00010; Millipore, Burlington, USA). Membranes were blocked with 5% non-fat milk and probed with primary antibody: Anti-MEF2A (#9736, CST, Danvers, USA); Anti-SIRT1 (#9475, CST, Danvers, USA); Anti-PGC-1α (#2178, CST, Danvers, USA); Loading controls: β-actin (#4970, CST, Danvers, USA) and GAPDH (#5174, CST, Danvers, USA). Band quantification used Image J with background subtraction.

## Statistical Analysis

In vitro experiments were independently repeated ≥ 3 times with data expressed as mean ± SEM. Mouse studies included ≥ 5 animals per group (biological replicates n ≥ 5). Normality was assessed using Shapiro-Wilk test. Parametric data were analyzed by two-tailed Student’s t-test; non-parametric data used Mann-Whitney U test. All analyses were performed in GraphPad Prism 8.0.1 with significance threshold set at P <0.05.

## Results

### MEF2A Deficiency Exacerbates Oxidative Stress via Mitochondrial Dysregulation

Building on our prior findings that MEF2A overexpression mitigates H_2_O_2_-induced endothelial senescence [27], we first interrogated its role in redox homeostasis. HUVECs exposed to 200 μM H_2_O_2_ for 2 hours exhibited an average of 37.4% reduction in viability (Figure 1A), paralleled by a marked downregulation of MEF2A protein (an average of 70% decrease; Figure 1B). Strikingly, MEF2A silencing recapitulated the oxidative phenotype of H_2_O_2_-treated cells: intracellular ROS surged by an average of 4.69-fold (Figure 1C), while mitochondrial ROS increased by an average of 67% (Figure 1D). Redox imbalance was further evidenced by a 46.6% reduction in GSH/GSSG ratio (Figure 1E), a 34% increase of NADH/NAD^+^ ratio (Figure 1F), and a 63.7% reduction in mitochondrial membrane potential (Figure 1G). With the silencing of MEF2A, there was a sharp decrease in mitochondrial membrane potential and a significant increase in mitochondrial ROS and NADH/NAD^+^ ratio, indicating that the strong oxidative stress caused by MEF2A deficiency is mainly due to the collapse of mitochondrial function.

**Figure 1.**
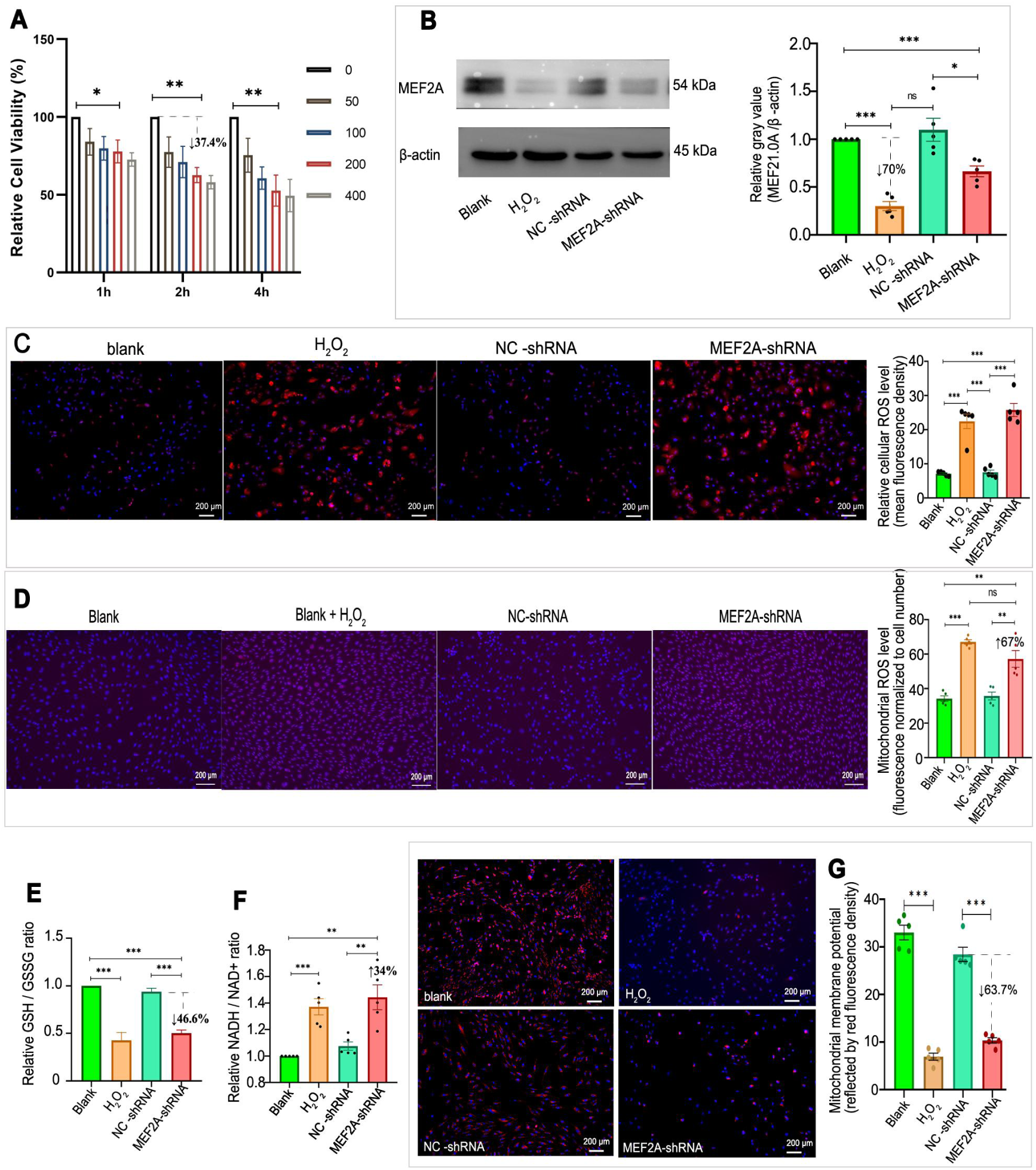
Modulation of MEF2A expression affects oxidative stress parameters in a cell model. (A) The bar graphs illustrate the viability of HUVECs upon exposure to different concentrations (μM) of H₂O₂ for different time (hours) in comparison to the control group. (B) Western blot analysis was performed to detect the expression levels of MEF2A. The blots show the bands corresponding to MEF2A (54 kDa) and β-actin (45 kDa) as a loading control. The right panel depicts the grayscale quantification of the MEF2A protein levels relative to the blank group after normalization with β-actin. (C) Fluorescence microscopy images of HUVECs stained with dihydroethidium were obtained to evaluate the levels of reactive oxygen species (ROS). Increased red fluorescence indicates higher ROS levels. The bar graphs show the mean relative fluorescence intensity normalized to cell number for each group. (D) Fluorescence microscopy images of HUVECs stained with OM08 (BestBio, equivalent to Mitosox™ Red from thermo) probe were obtained to evaluate the levels of mitochondrial ROS (mt-ROS). Increased red fluorescence indicates higher mt-ROS levels. The bar graphs display the mt-ROS normalized to cell number in different groups. (E) Bar graphs presenting the GSH/GSSG ratio relative to the control group. (F) The bar graphs illustrate the NADH/NAD⁺ ratio relative to the control group. (G) Fluorescence microscopy images of HUVECs stained with Tetramethylrhodamine ethyl ester (TMRE) were used to assess the mitochondrial membrane potential. The bar graph shows the fluorescence intensity normalized to the cell number. Data for all bar plots are represented as the mean ± standard error from multiple independent experiments (n = 3 - 5). Student’s t-test was used to compare the significance of differences between two groups. *, *P* < 0.05; **, *P* < 0.01; ***, *P* < 0.001.

### MEF2A Orchestrates the SIRT1/PGC-1**α** Axis to Sustain Redox Homeostasis

Given the established role of SIRT1/PGC-1α in mitochondrial integrity[29], we hypothesized this pathway mediates MEF2A’s antioxidant effects. Silencing MEF2A reduced SIRT1 and PGC-1α protein levels by an average of 34.4% and 38.8%, respectively (Figure 2A). Notably, SIRT1 knockdown recapitulated MEF2A deficiency, elevating cellular ROS (2.4-fold; Figure 2B) and mitochondrial ROS (Figure S1), disrupting NADH/NAD^+^ balance (31.8% increase in NADH/NAD^+^ ratio; Figure 2C), reducing the GSH/GSSG ratio by 46% (Figure 2D), and reducing mitochondrial membrane potential by 53.6% (Figure 2E). Rescue experiments confirmed functional interdependence: SIRT1 overexpression ameliorated MEF2A knockdown-induced PGC-1α suppression (Figure 2A) and restored all cellular phenotypes— suppressing cellular ROS (Figure 2B) and mitochondrial ROS (Figure S1), normalizing NADH/NAD^+^ ratios (Figure 2C), increasing GSH/GSSG ratios (Figure 2D), and rescuing mitochondrial membrane potential (Figure 2E). These data establish MEF2A as a critical upstream regulator functionally sustaining the SIRT1/PGC-1α axis, which is essential for redox and mitochondrial homeostasis.

**Figure 2.**
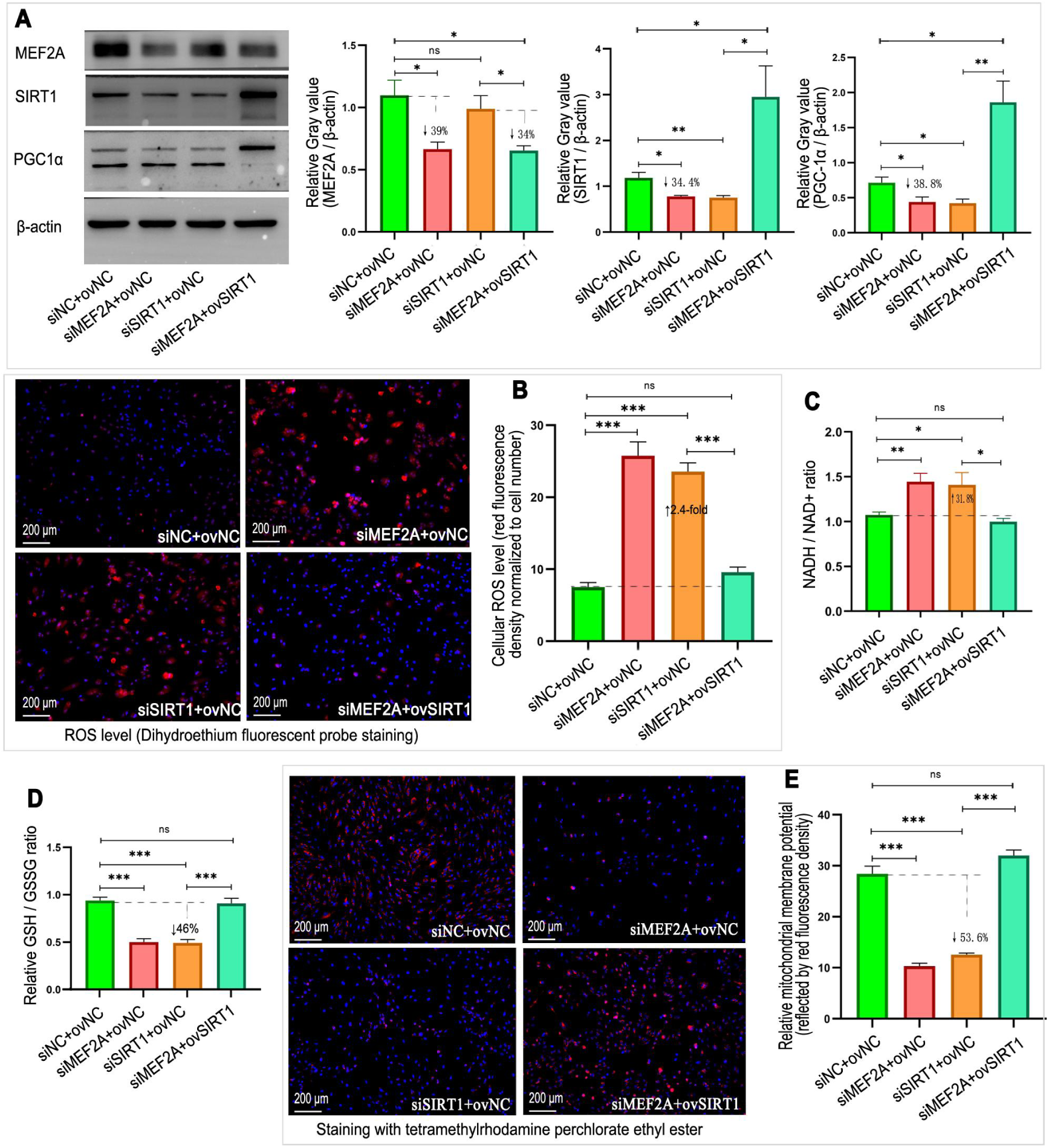
The Influence of MEF2A Silencing on SIRT1/PGC-1**α** Expression and Cellular Oxidative Stress (A) Western blot analysis was carried out, and the grayscale value plots relative to β-actin were presented. (B) Fluorescence microscopy images of HUVECs under different interventions involving MEF2A or SIRT1 were obtained. The enhanced red fluorescence indicates an increase in ROS levels. Corresponding bar graphs were used to depict the relative average ROS levels in each group, providing a quantitative comparison. (C) Bar graphs were constructed to show the mean relative NADH/NAD^+^ ratio for each group. (D) The mean relative GSH/GSSG ratio per group was presented in bar graphs. (E) Fluorescence microscopy images of HUVECs stained with TMRE for the detection of mitochondrial membrane potential were shown. Accompanying bar charts of the mean relative fluorescence intensity per group were also provided to facilitate comparison among different groups. Error bars represent the standard error derived from multiple independent experiments (n = 3 - 5). Student’s t-test was used to compare the significance of differences between two groups. si-MEF2A represents MEF2A silencing, si-SIRT1 represents SIRT1 silencing, si-NC is the siRNA negative control, and ov-NC is the overexpression plasmid negative control. *, *P <* 0.05; **, *P <* 0.01; ***, *P <* 0.001.

### Activating the expression of MEF2A may be a highly effective new method for antioxidant therapy through SIRT1/PGC-1**α** axis

Following the overexpression of MEF2A in HUVECs, there was a significant upregulation of SIRT1 (Figure 3 A, C) and PGC-1α (Figure 3A, D) expression. Overexpressing SIRT1 did not affect the expression of MEF2A (Figure 3A, B). When SIRT1 was silenced simultaneously with the overexpression of MEF2A, the expression level of SIRT1 showed no significant change compared to the negative control group (Figure 3A, C). Overexpressing MEF2A produced the same antioxidative stress effects as overexpressing SIRT1, including a significant reduction in ROS levels (Figure 3E), mitochondrial ROS levels (Figure 3F), and the NADH/NAD^+^ ratio (Figure 3G) in H_2_O_2_-treated cells, a significant increase in the GSH/GSSG ratio (Figure 3H), and a significant increase in mitochondrial membrane potential (Figure 3I). When cells were treated with H_2_O_2_ while SIRT1 was silenced simultaneously with the overexpression of MEF2A, the cellular effects were consistent with those of the negative control group (Figure 3E-I). This result indicates that overexpression of MEF2A and overexpression of SIRT1 have almost identical antioxidant effects, and overexpression of MEF2A achieves its antioxidant function by upregulating the expression of SIRT1.

**Figure 3.**
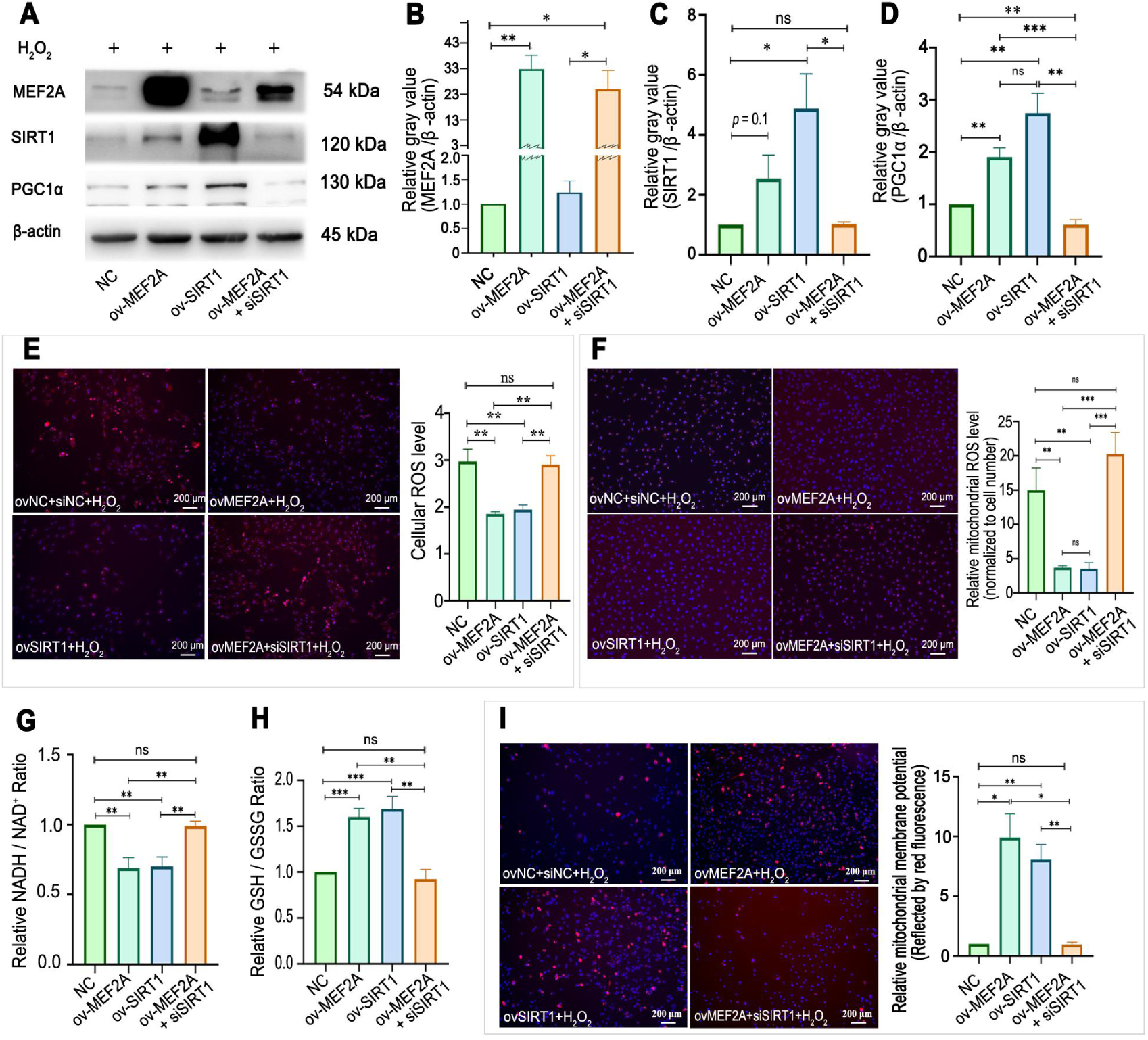
Effects of MEF2A overexpression on oxidative stress and the SIRT1/PGC-1**α** pathway. (A) Western blot bands of various genes (β-actin band as the load control). The bars show the relative expression levels of MEF2A (B), SIRT1 (C), and PGC-1α (D) represented with grayscale value normalized to β-actin. (E) Fluorescence microscopy images of HUVECs stained with dihydroethidium were obtained to evaluate the levels of the cellular ROS levels. Increased red fluorescence indicates higher ROS levels, with bar graphs representing relative average ROS levels per group. (F) Fluorescence microscopy images of HUVECs stained with OM08 probe were obtained to evaluate the levels of mitochondrial ROS (mt-ROS). Increased red fluorescence indicates higher mt-ROS levels. The bar graphs display the mt-ROS normalized to cell number in different groups. (G) Bar graphs illustrating the mean relative NADH/NAD^+^ ratio for each group. (H) Bar graphs depicting the mean relative GSH/GSSG ratio per group. (I) Fluorescence microscopy images of HUVECs stained with TMRE to assess mitochondrial membrane potential, with bar charts showing mean relative fluorescence intensity per group. DAPI (4’,6-diamidino-2-phenylindole) stained the cell nucleus blue. Data for all bar plots are represented as the mean ± standard error from multiple independent experiments (n = 3-5). Student’s t-test was used to compare the significance of differences between two groups. ov-MEF2A, MEF2A overexpression; ov-SIRT1, SIRT1 overexpression; NC, negative control; *, *P <* 0.05; **, *P <* 0.01; ***, *P <* 0.001.

### MEF2A Directly Transactivates SIRT1 via Conserved Promoter Elements

Although previous literature has suggested that MEF2A regulates the expression of SIRT1, the molecular mechanism by which MEF2A regulates SIRT1 expression is not fully understood. Here, mechanistic dissection revealed two evolutionarily conserved MEF2A binding motifs in the SIRT1 promoter: motif-1 (-393/-378 bp) and motif-2 (-904/-890 bp) (Figure 4A). Luciferase assays demonstrated that wild-type SIRT1 promoter activity exceeded mutant control by 43% under basal conditions (Figure 4B) and by 77% upon MEF2A overexpression (Figure 4C). Furthermore, co-expression of MEF2A enhanced wild-type SIRT1 promoter activity by 73% compared to basal conditions (Figure 4D). Chromatin immunoprecipitation (ChIP) validated direct binding: anti-MEF2A enriched SIRT1 promoter fragments by >10-fold vs. IgG controls (Figure 4E-G). This transcriptional regulation is unidirectional, as SIRT1 knockdown did not alter MEF2A levels (Figure 2A), cementing MEF2A as a primary driver of SIRT1 expression.

**Figure 4.**
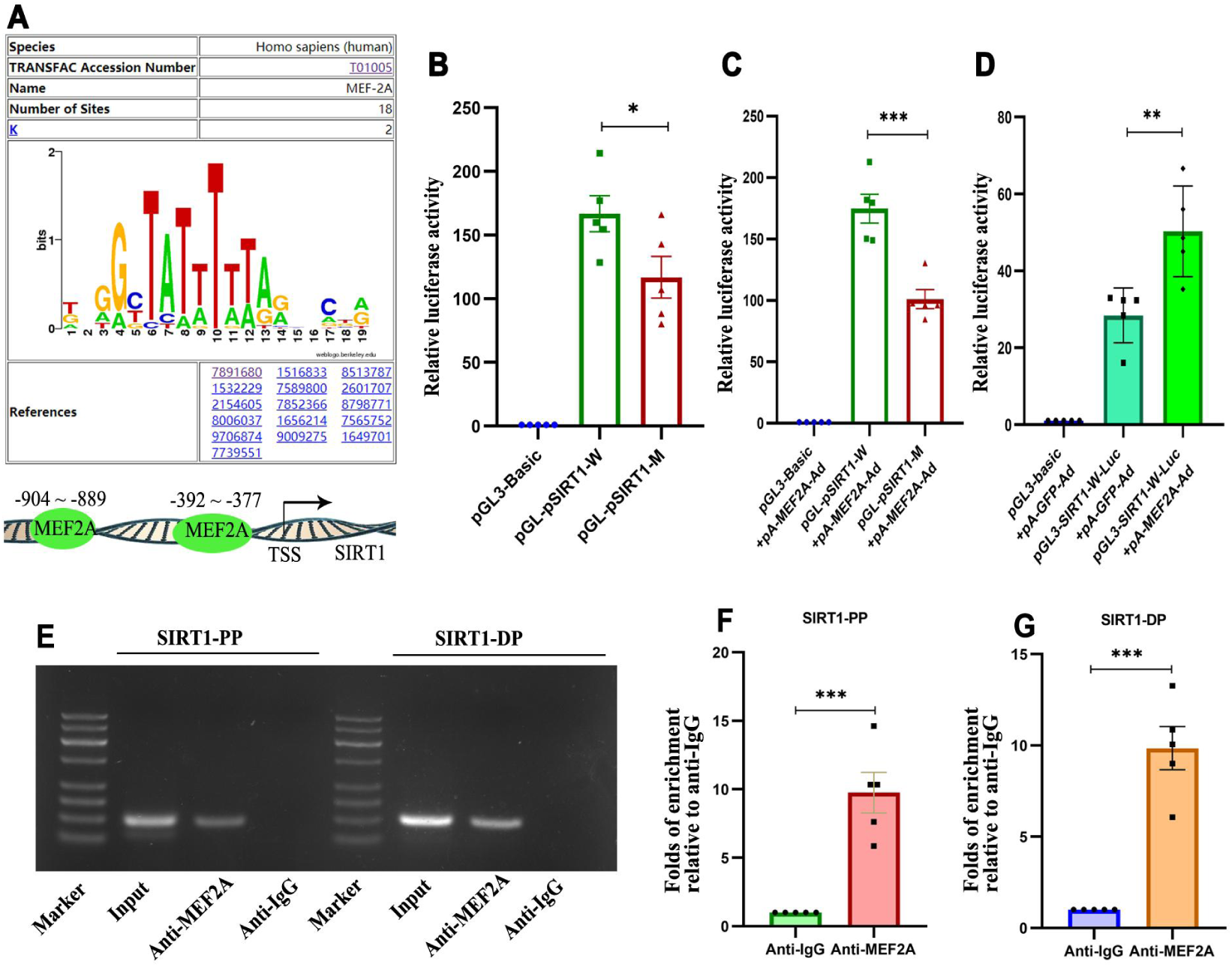
Prediction and validation of MEF2A binding sites in the SIRT1 promoter region. **(A)** Characteristic sequence of MEF2A binding sites and a schematic of MEF2A binding in the SIRT1 promoter region, as predicted by TF-Search2.0. **(B)** The relative luciferase activity was calculated by dividing the normalized value of each group in each experiment (firefly luciferase activity / Renilla luciferase activity) by the normalized value of the control group (pGL3-basic). **(C)** The relative luciferase activity was calculated by dividing the normalized value of each group in each experiment (firefly luciferase activity / Renilla luciferase activity) by the normalized value of the control group (pGL3-basic + pA-MEF2A-Ad). **(D)** The relative luciferase activity was calculated by dividing the normalized value of each group in each experiment (firefly luciferase activity / Renilla luciferase activity) by the normalized value of the control group (pGL3-basic + pA-GFP-Ad). **(E)** Gel electrophoresis results of PCR products amplified from DNA fragments pulled down by different antibodies in ChIP. **(F and G)** Quantitative analysis of SIRT1 proximal and distal promoter DNA fragments pulled down by anti-MEF2A using qPCR. Enrichment fold was calculated by using the formula 2^-△Ct^ (△Ct = Ct_(anti-MEF2A)_ - Ct_(anti-IgG)_). The bar charts present the data as mean ± SEM. All experiments were performed in at least three independent replicates. Use Student’s t-test to compare the significance of differences between two groups. *, *P <* 0.05; **, *P <* 0.01; ***, *P <* 0.001. pGL3-basic, luciferase plasmid without promoter; pGL-pSIRT1-W, luciferase plasmid with wild-type SIRT1 promoter; pGL-pSIRT1-M, luciferase plasmid with SIRT1 promoter mutated at predicted MEF2A binding sites. pA-MEF2A-Ad, adenovirus expressing MEF2A. SIRT1-PP, DNA fragment near MEF2A binding sites in the proximal SIRT1 promoter; SIRT1-DP, DNA fragment near MEF2A binding sites in the distal SIRT1 promoter.

Collectively, these in vitro experimental results suggest that unlocking the MEF2A-SIRT1 master switch may be a novel paradigm for targeted antioxidant therapy.

### Endothelial-Specific MEF2A Knockdown Recapitulates Vascular Oxidative Pathology In Vivo

To validate the antioxidant role of MEF2A in vascular tissues in vivo, we generated endothelial-specific Mef2a knockdown mice (C57BL/6J background) and maintained them on a high-fat diet (HFD) for four months to assess vascular pathology and gene expression alterations.

Fluorescence imaging of fresh carotid artery sections revealed robust GFP expression throughout the vascular wall (Figure 5A), confirming successful AAV1 transduction and transgene expression in vascular tissues. Faint green fluorescence was also observed in frozen sections of liver, kidney, spleen, and lung tissues (data not shown). While this signal may originate from abundant microvasculature within these organs, the endothelial specificity of MEF2A expression was confirmed through tissue-specific knockdown efficacy analysis: Western blot demonstrated unchanged MEF2A levels in liver (Figure 5B) and heart tissues (Figure 5C), but significantly decreased in aortic tissues compared to controls (Figure 5D). Following 16 weeks of HFD feeding, Mef2a knockdown mice exhibited increased body weight gain compared to controls (Figure 5E), while plasma lipid profiles (total cholesterol, HDL-C, LDL-C) and blood glucose levels remained unchanged (Figure 5F-I). Notably, circulating triglycerides were markedly elevated in the knockdown group (Figure 5J).

**Figure 5.**
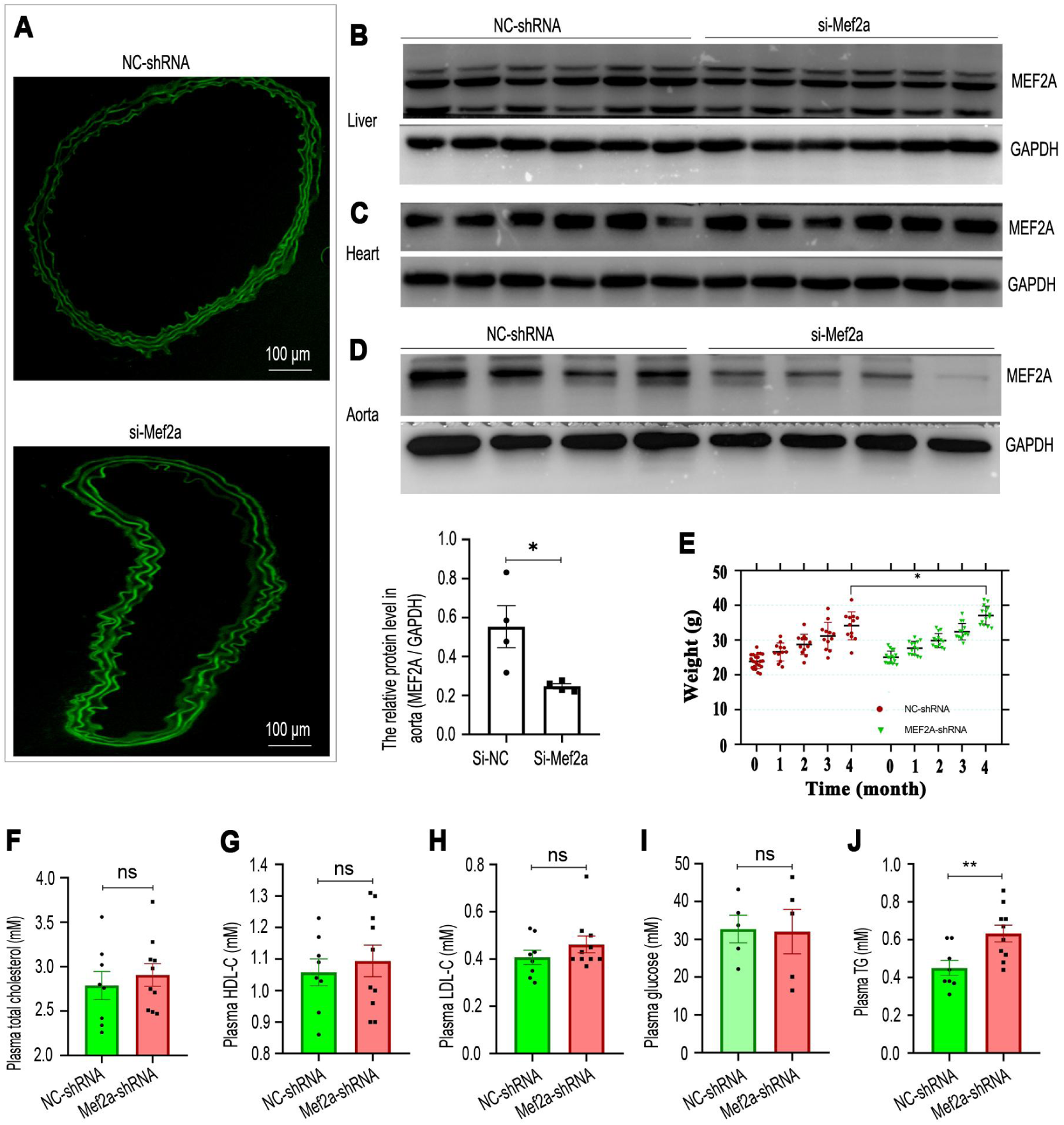
Targeted knockdown of MEF2A in mouse blood vessel and its physiological effects. **(A)** Fluorescence microscopy images of carotid frozen sections showing the expression of GFP in the vascular ring. The upper image represents the NC - shRNA group, and the nether image represents the si-Mef2a (Mef2a knockdown) group, with a scale bar of 100 μm. **(B - D)** Western blots were used to assess the expression levels of MEF2A in different tissues. **(B)** MEF2A expression in the mouse liver (n = 6 biological replicates), with GAPDH as a loading control. **(C)** MEF2A expression in the mouse heart (n = 6 biological replicates), with GAPDH as a loading control. **(D)** MEF2A expression in the mouse aorta (n = 4 biological replicates), with GAPDH as a loading control The bar chart with scatter plot indicate the relative protein level of MEF2A quantified by grayscale value analysis. Data are presented as mean ± SEM. The biological replicates of aortic tissue here are only 4, which is less than those of liver and heart tissue, mainly because aortic tissue needs to be used for more detection items, such as Oil Red O staining, mRNA extraction, etc. The 4 aortic tissues here are from different biological replicates compared to the 5 aortic tissues in Figure 7 **(E)** Comparison of mouse weights between the NC and si-MEF2A groups at various time points during a high - lipid diet. The data is represented by a scatter plot with median and interquartile range. (F - J) The bar charts with scatter plot indicate the relative levels of blood lipid and glucose levels after 4 months of feeding between the NC and si-MEF2A groups. Data are presented as mean ± SEM. (F) Plasma total cholesterol levels. (G) Plasma high density lipoprotein cholesterol (HDL-C) levels. (H) Plasma low density lipoprotein cholesterol (LDL-C) levels. (I) Plasma glucose levels. (J) Plasma triglycerides (TG) levels. Use Student’s t-test or Mann Whiteley U test to compare the significance of differences between two groups. ns, not statistically significant; *, *P* < 0.05; **, *P* < 0.01

Mef2a silencing induced pronounced vascular pathology: 1) Accelerated atherogenesis: En face Oil Red O staining revealed significantly increased lipid deposition in aortic arches (Figure 6A); 2) Systemic redox imbalance: Plasma GSH/GSSG ratio was substantially reduced (Figure 6B) and tissue ROS was significantly increased (Figure 6C); 3) DNA oxidative damage: Vascular 8-hydroxy-2’-deoxyguanosine (8-OHDG) levels, a biomarker of ROS-mediated DNA damage [30], were elevated (Figure 6D).

**Figure 6.**
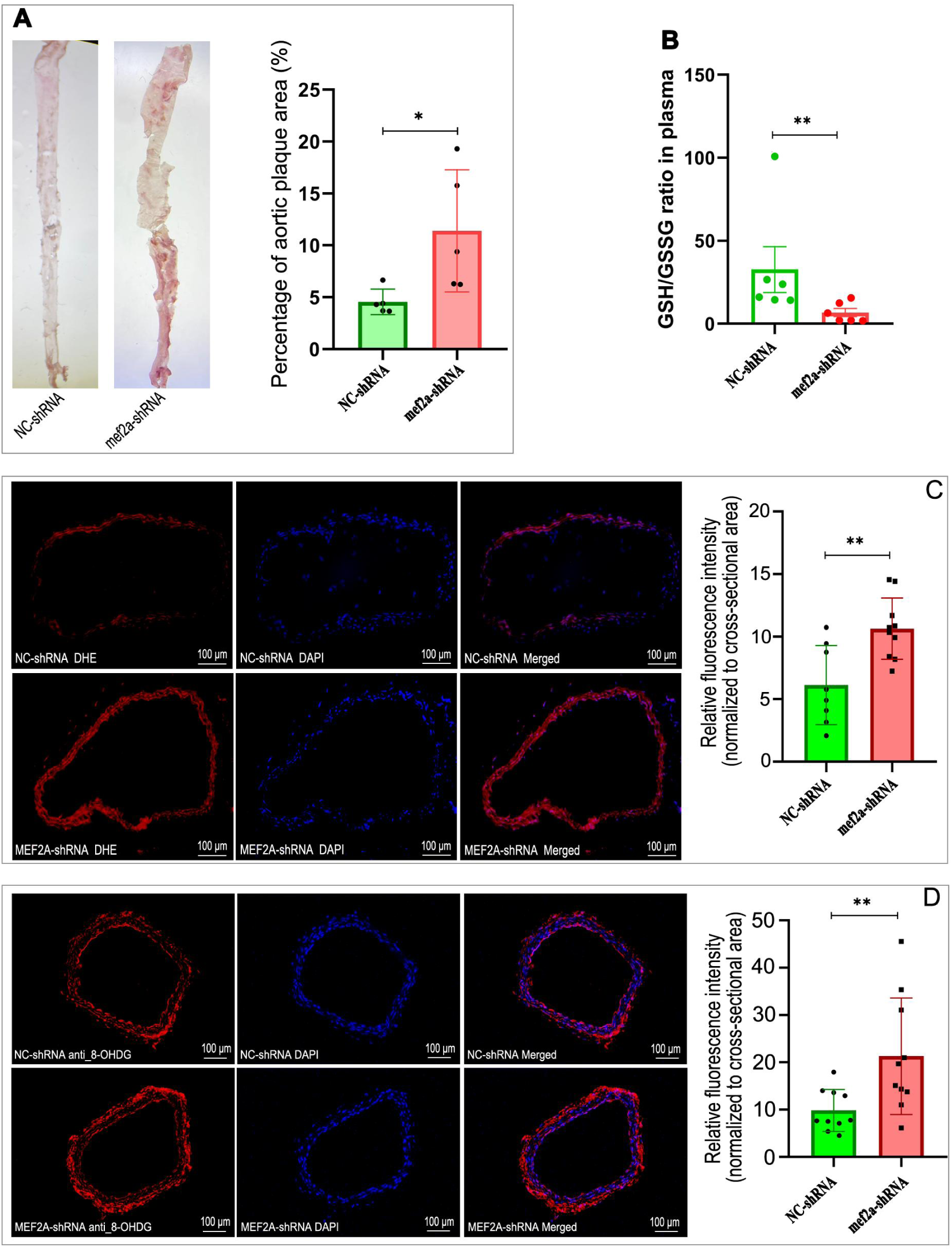
Pathological Alterations in Mice with Specific MEF2A Knockdown in Blood Vessel. (A) En-face staining of the aorta with Oil Red O. The combined bar-scatter plot shows the mean percentage of aortic plaque area. (B) The combined bar-scatter plot shows the mean values of plasma GSH/GSSG ratio for each group. (C) Fluorescence microscopy images of frozen carotid sections were obtained after staining with a dihydroethidium (DHE) probe to evaluate the levels of reactive oxygen species (ROS). Red fluorescence indicates a high level of ROS. DAPI (4’,6-diamidino-2-phenylindole) stained the cell nucleus blue. The combined bar-scatter plot shows the mean relative red fluorescence intensity normalized to the vascular ring area for each experimental group. (D) Fluorescence microscopy images of paraffin-embedded carotid sections incubated with antibody against 8-hydroxy-2′-deoxyguanosine (8-OHDG). Red fluorescence indicates generation of 8-OHDG that is a marker of DNA damage. DAPI stained the cell nucleus blue. The combined bar-scatter plot shows the mean relative red fluorescence intensity normalized to the vascular ring area for each experimental group. Student’s t-test was used to compare the significance of differences between two groups. The error bar represents S.E.M., and significant differences are indicated with * (*P* < 0.05), and ** (*P* < 0.01).

Western blot and quantitative PCR (qPCR) analyses confirmed coordinated downregulation of both Sirt1 and Pgc1a in aortic tissues from Mef2a knockdown mice, with significant reductions observed at protein (Figure 7A-F) and mRNA levels (Figure 7G-I). The results shown in Figure 7A and Figure 5D were derived from different individuals within the corresponding groups of the same cohort of Mef2a knockdown mice and their negative control mice. This in vivo validation aligns with our in vitro findings, demonstrating that MEF2A deficiency promotes vascular oxidative stress and DNA damage through suppression of the SIRT1/PGC-1α axis.

**Figure 7.**
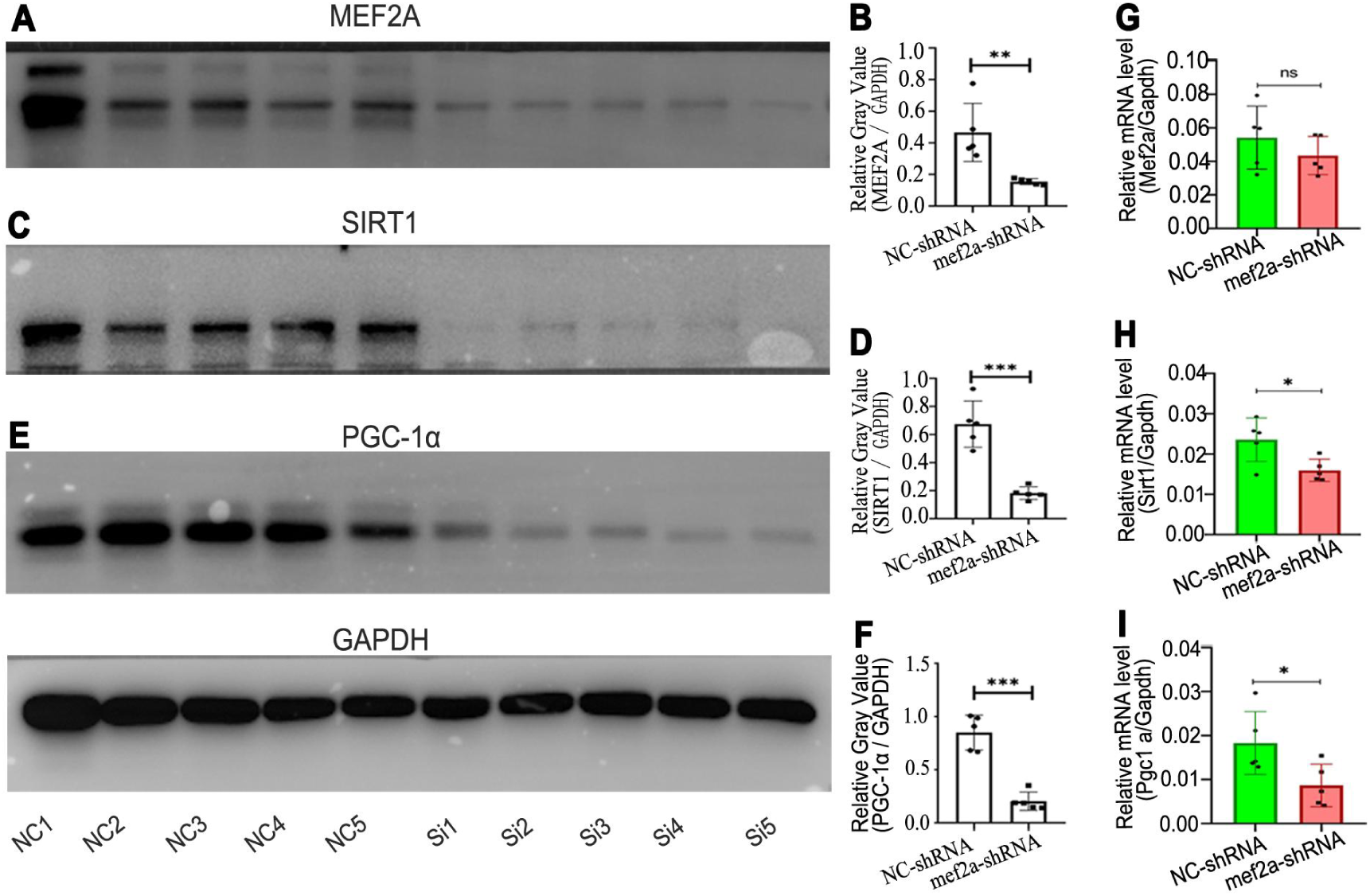
Western Blot and Quantitative Real-time PCR (qPCR) Analyses for Gene Expression in Mouse Aorta. **A, C, E**: Western blot analyses were performed for MEF2A (**A**), SIRT1 (**C**), and PGC-1α (**E**) proteins using respective antibodies. Lanes NC1 - NC5 and Si1 - Si5 respectively represent different negative control (NC) and Mef2a knockdown (Si) samples from 5 biological replicates, with GAPDH serving as a loading control. **B, D, F**: The relative protein levels of MEF2A (**B**), SIRT1 (**D**), and PGC-1α (**F**) were quantified by grayscale value analysis. The combined bar-scatter plot present the data as mean ± SEM. **G, H, I**: qRT-PCR was used to analyze the mRNA levels of *Mef2a* (**G**), *Sirt1* (**H**), and *Pgc1a* (**I**). The combined bar-scatter plot show the mRNA levels relative to that of internal reference gene (Gapdh). Data are presented as mean ± SEM. The mark’ns’ indicates no statistically significant difference. Student’s t-test was used to compare the significance of differences between two groups. The symbols *, **, and *** indicate statistically significant differences between NC and Mef2a knockdown groups at *P* < 0.05, *P* < 0.01, and *P* < 0.001, respectively.

## Discussion

Our study establishes MEF2A as a central transcriptional orchestrator of vascular redox homeostasis, functioning through direct governance of the SIRT1/PGC-1α axis. While oxidative stress is universally acknowledged as a pivotal mechanism underlying endothelial dysfunction and atherosclerotic progression[31–33], contemporary therapeutic strategies remain ineffective in targeting its fundamental molecular drivers[34]. The present work demonstrated that MEF2A deficiency precipitated catastrophic redox imbalance characterized by uncontrolled reactive oxygen species (ROS) generation, glutathione system collapse, and mitochondrial dysfunction—pathological hallmarks mirroring advanced atherosclerosis[35]. Crucially, MEF2A overexpression not only reversed these abnormalities but also restored physiological redox signaling, establishing its indispensable role in vascular antioxidant defense.

The mechanistic hierarchy of MEF2A-mediated protection operates through coordinated regulation of mitochondrial quality control systems. Chromatin occupancy studies confirmed MEF2A binding to evolutionarily conserved cis-regulatory elements within the SIRT1 promoter, driving transcriptional activation essential for maintaining SIRT1 expression. This genomic control synergizes with post-translational modulation of nicotinamide adenine dinucleotide (NAD^+^) metabolism, wherein MEF2A signaling stabilizes cellular NAD^+^ pools required for SIRT1 deacetylase activity[36–38]. Complementing these mechanisms, MEF2A induces PGC-1α expression to optimize mitochondrial biogenesis while suppressing electron transport chain-derived ROS leakage[12,13,39,40]. Collectively, these findings elucidate the critical role of MEF2A in maintaining vascular redox homeostasis through direct transcriptional activation of SIRT1, thereby modulating the SIRT1/PGC-1α axis. This finding not only advances our understanding of vascular oxidative stress regulation but also identifies a potential therapeutic target for oxidative cardiovascular diseases.

Oxidative stress, characterized by an imbalance between reactive oxygen species (ROS) production and antioxidant defenses, is a key contributor to cardiovascular pathologies such as atherosclerosis and hypertension. In atherosclerosis, redox imbalance promotes lipid peroxidation, particularly of low-density lipoprotein (LDL), leading to the formation of oxidized LDL (oxLDL) and subsequent endothelial dysfunction, inflammation, and foam cell formation—hallmarks of plaque development[41,42]. Similarly, in hypertension, metabolic disruptions like glucose-6-phosphate dehydrogenase (G6PD) deficiency induce a metabolic switch that exacerbates oxidative stress, driving pulmonary vascular remodeling and hypertension[43]. These mechanisms underscore the therapeutic potential of targeting redox imbalance in cardiovascular diseases.

The SIRT1/PGC-1α pathway is central to mitochondrial quality control and antioxidant defense, making it a focal point in cardiovascular research. SIRT1, a NAD^+^-dependent deacetylase, deacetylates and activates PGC-1α, a master regulator of mitochondrial biogenesis and ROS detoxification[29,44]. This axis is critical for preserving endothelial function and mitigating oxidative damage [12,37]. Recent studies highlight its therapeutic relevance: compounds like urolithin A and resveratrol activate SIRT1/PGC-1α to ameliorate mitochondrial dysfunction and oxidative stress in models of Parkinson’s disease and diabetic nephropathy [14,22,45]. Deeper mechanistic exploration of this pathway, including its upstream regulators (e.g., MEF2A) and downstream effectors (e.g., mitochondrial dynamics), could unveil novel strategies to counteract oxidative cardiovascular diseases.

These mechanistic insights carry profound therapeutic implications that challenge conventional antioxidant paradigms. Current clinical failures in redox-targeted interventions stem from two fundamental limitations: nonspecific ROS scavenging that disrupts physiological signaling, and inadequate attention to transcriptional networks governing endogenous defense systems[42,46,47]. Our results position MEF2A activation as a precision therapeutic strategy that simultaneously enhances mitochondrial integrity upon upregulating SIRT1/PGC-1α, bolsters endogenous antioxidant capacity, and preserves redox-sensitive vascular homeostasis. The spatial specificity of this regulation is particularly noteworthy—endothelial-restricted MEF2A knockdown in murine models induced severe vascular oxidative pathology while sparing cardiac and hepatic tissues, suggesting cell-type-specific delivery systems could maximize therapeutic efficacy while minimizing off-target effects.

Reconciling MEF2A’s dual roles in vascular protection and disease pathogenesis requires understanding its compartment-specific biology. Although MEF2A is the core transcription factor for myocardial cell hypertrophy[48], our findings reveal a protective endothelial role through nuclear SIRT1/PGC-1α activation. This apparent paradox resolves through subcellular localization dynamics: nuclear MEF2A in vascular endothelium promotes redox homeostasis, whereas cytoplasmic MEF2A in cardiomyocytes may drive pathological hypertrophy through calcineurin-dependent mechanisms[49]. Such tissue-specific functionality underscores the importance of developing targeted delivery platforms that exploit MEF2A’s protective capacity without activating deleterious pathways in other cell types.

The molecular mechanisms by which MEF2A regulates redox balance are likely multifaceted. Although our study observed that MEF2A exerts anti-oxidative stress effects primarily by directly regulating SIRT1 to upregulate PGC-1α, there are also reports suggesting that MEF2A can directly modulate PGC-1α to promote mitochondrial biogenesis in gastric cancer cells [50]. It may even enhance mitochondrial ROS clearance by inhibiting KEAP1 to stabilize NRF2, thereby exerting anti-oxidative effects [50]. These findings indicate that MEF2A may employ multiple parallel pathways to exert its anti-oxidative functions. The predominant pathway may vary depending on cell type and the specific microenvironment under physiological or pathological conditions. Notably, our study observed that MEF2A not only promotes SIRT1 expression but also may enhance SIRT1 activity by increasing NAD^+^ levels. Collectively, these studies demonstrate that MEF2A activates the antioxidant system through diverse mechanisms. In drug screening, adopting MEF2A upregulation as a criterion for selecting anti-oxidative agents might prove to be a more effective strategy. Indeed, the protective effect of the antioxidant resveratrol on vascular endothelial cells depends on the up regulation of MEF2A expression [22].

The evolutionary conservation of MEF2 family proteins offers additional insights into its therapeutic potential. From Drosophila to mammals, MEF2 homologs have maintained dual roles in stress adaptation and metabolic regulation—an ancient functional duality that modern medicine might exploit[51,52]. Our finding that MEF2A coordinates mitochondrial renewal through SIRT1/PGC-1α parallels its role in maintaining muscle mitochondrial networks during aging, suggesting conserved mechanisms for combating oxidative decline across tissues.

Despite these advances, several critical questions must guide future investigation. First, the cell-type-specific effects of MEF2A modulation in complex vascular ecosystems remain incompletely characterized[15]. While our work establishes endothelial MEF2A as a master redox regulator, its potential roles in vascular smooth muscle cell phenotypic switching or macrophage polarization during atherogenesis warrant systematic exploration. Single-cell transcriptomic analyses of atherosclerotic plaques could delineate compartment-specific regulatory networks and identify cell populations most amenable to MEF2A-targeted interventions. Second, the integration of MEF2A signaling with biomechanical forces requires elucidation. Given MEF2A’s established mechanosensitivity and the well-documented atheroprotective effects of laminar shear stress[53,54], investigating potential synergy between hemodynamic forces and MEF2A activation could inform both pharmacological strategies and device-based therapies. Third, the functional speculation of MEF2A in atherosclerosis relies heavily on murine models; however, it is crucial to note the substantial biological differences between human and murine vasculature. Notably, murine atherosclerosis models lack key human-specific pathological features, such as fibrous cap composition and plaque calcification dynamics[55]. These limitations necessitate translational validation in larger animal models or human vascular systems to confirm clinical relevance. Fourth, while our ChIP assay provides evidence for MEF2A’s direct binding to the predicted cis-elements in the promoter of SIRT1, the assay regions being limited to the predicted binding sites may overlook potential indirect interactions between MEF2A and the lateral cis-elements. In order to obtain compelling evidence for the direct binding of proteins to specific DNA regions, the method used by Xie B., et al.[56] to decipher the binding sites of H3K18la in the LCN2 promoter is worthy of recommendation.

## Conclusion

This study establishes MEF2A as a central regulator of vascular redox homeostasis through coordinated activation of the SIRT1/PGC-1α axis and NAD^+^ stabilization.

## List of abbreviations

MEF2A: Myocyte Enhancer Factor 2A
ROS: Reactive Oxygen Species
HUVEC: Human Umbilical Vein Endothelial Cell
VECs: Vascular Endothelial Cell
8-OHDG: 8-Hydroxy-2’-Deoxyguanosine
9-GSH: Glutathione
GSSG: Glutathione Disulfide
NAD^+^: Oxidized Nicotinamide Adenine Dinucleotide
NADH: Nicotinamide Adenine Dinucleotide
SIRT1: Silent Information Regulator 1
PGC-1α: Peroxisome Proliferator-Activated Receptor γ Co-Activator 1α
H_2_O_2_: Hydrogen Peroxide
ChIP: Chromatin Immunoprecipitation
CVD: Cardiovascular Diseases
AAV1: Type 1 Adeno-Associated Virus
shRNA: Short-Hairpin RNA

## Acknowledgments

We would like to express our gratitude to Yang Ji, Yichao Deng, Huanzhen Chen, and Changnong Chen for their assistance in the experimental methods and process.

## Author contributions

Conceptualization and Writing – original draft, Benrong Liu; Investigation, Lei Fang and Chunxia Miao; Data curation, Lei Fang; Formal analysis, Chunxia Miao and Yujuan Xiong; Methodology, Xinyu Wen; Validation, Xiumiao Zheng; Writing – review & editing, Benrong Liu and Shi-Ming Liu; Funding acquisition, Benrong Liu and Shi-Ming Liu.

## Data availability

The experimental data generated and/or analyzed during this study are available from the corresponding author upon reasonable request.

## Declarations

The authors declare no competing interests.

## Supplementary Figures

**Figure S1.**
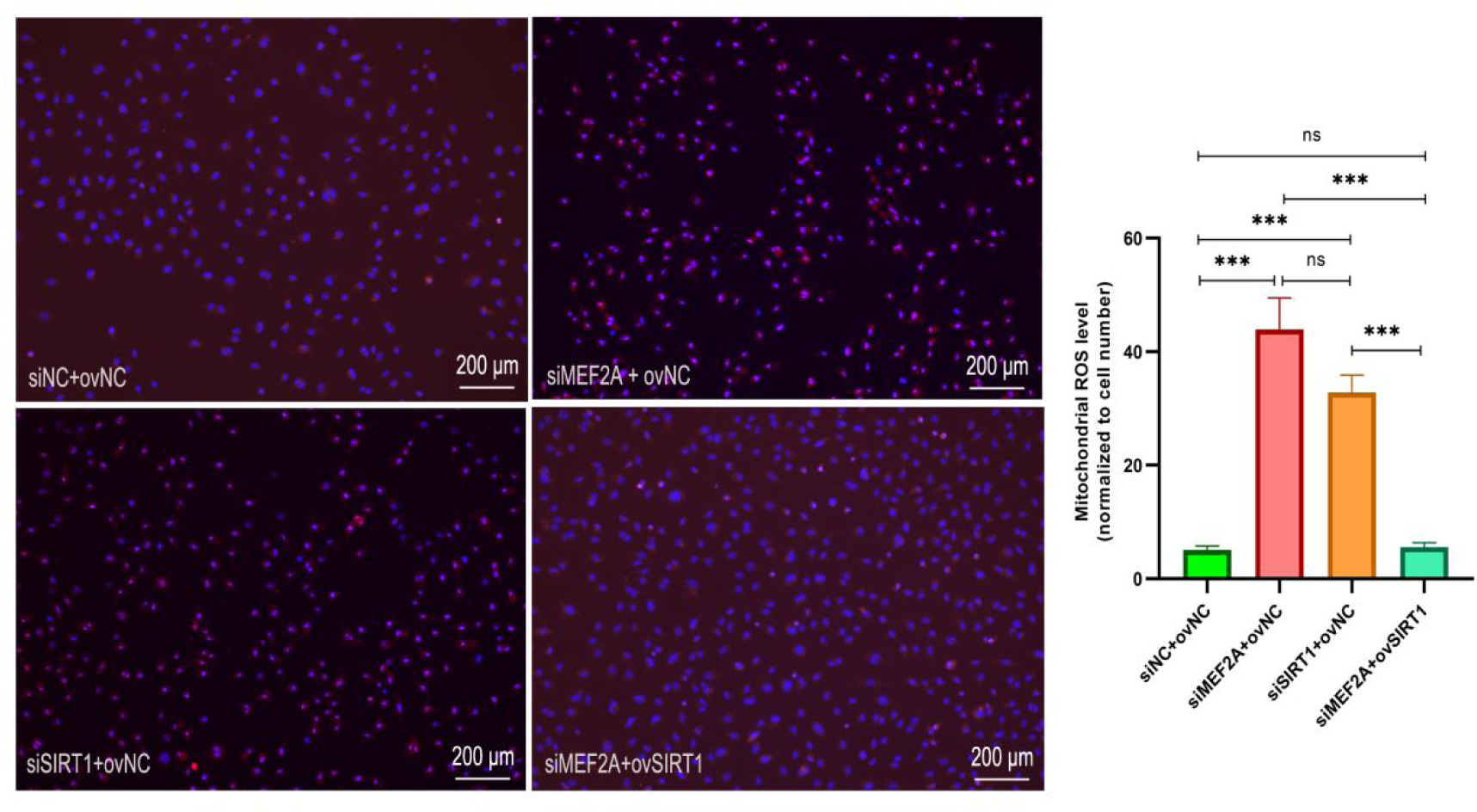
Mitochondrial ROS Levels in Different Experimental Groups after Staining with OM08 Probe. Fluorescence microscopy images of mitochondrial ROS staining using the OM08 (Bestbio) probe in different experimental groups are shown. DAPI (4’,6-diamidino-2-phenylindole) stained the cell nucleus blue, red fluorescence indicate high level of ROS. The groups include siNC + ovNC, siMEF2A + ovNC, siSIRT1 + ovNC, and siMEF2A + ovSIRT1. Each image has a scale bar of 200 μm. The right side of the figure presents a bar graph depicting the mean fluorescence intensity of mitochondrial ROS levels, normalized to the number of cells in each group. The error bars represent standard error. Statistical significance was determined using Student’s T test. ns indicates no significant difference, *** indicates *P* < 0.001.

## Supplementary Tables

**Table S1.**
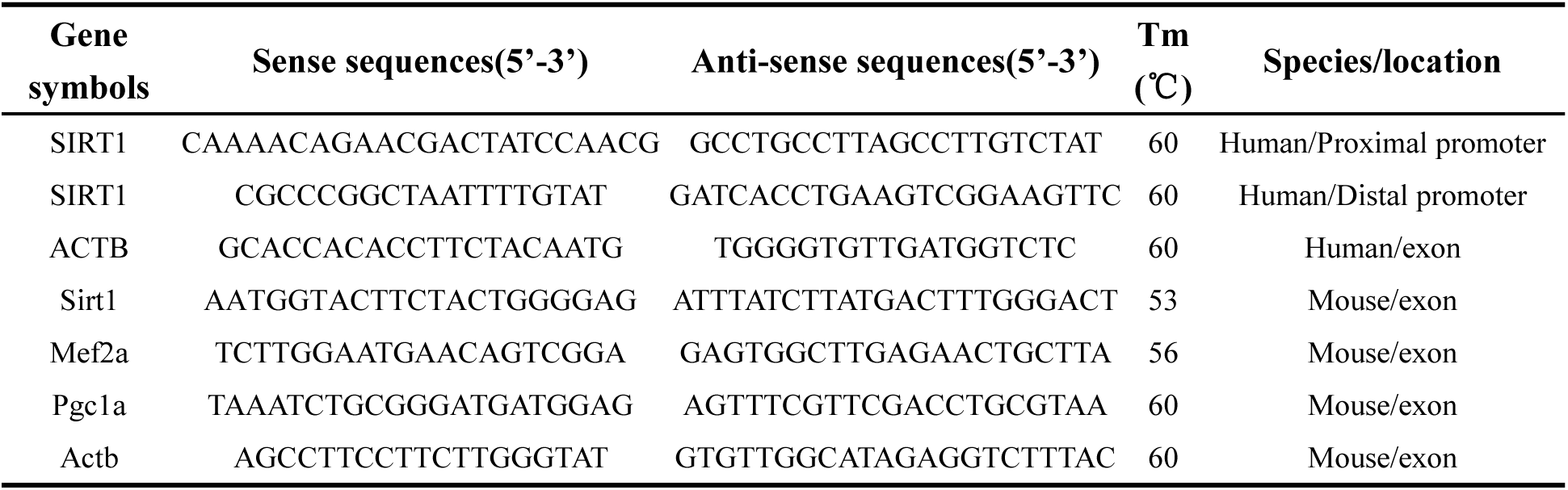
Prime sequences for quantitative real-time PCR and general PCR.

## References

1. Akiyama Y, Ivanov P. Oxidative Stress, Transfer RNA Metabolism, and Protein Synthesis. Antioxid Redox Signal 2024, 40: 715–735

2. Huang X, Zhou Y, Guo Y, Yan D, Sun P, Cao Y, Chen Y, et al. Selenium-Doped Copper Formate Nanozymes with Antisenescence and Oxidative Stress Reduction for Atherosclerosis Treatment. Nano Lett 2025, 25: 2662–2669

3. Zheng Y, Shao M, Zheng Y, Sun W, Qin S, Sun Z, Zhu L, et al. PPARs in atherosclerosis: The spatial and temporal features from mechanism to druggable targets. J Adv Res 2025, 69: 225–244

4. Griendling KK, Camargo LL, Rios FJ, Alves-Lopes R, Montezano AC, Touyz RM. Oxidative Stress and Hypertension. Circ Res 2021, 128: 993–1020

5. Kibel A, Lukinac AM, Dambic V, Juric I, Selthofer-Relatic K. Oxidative Stress in Ischemic Heart Disease. Oxid Med Cell Longev 2020, 2020: 6627144

6. Shaito A, Aramouni K, Assaf R, Parenti A, Orekhov A, Yazbi AE, Pintus G, et al. Oxidative Stress-Induced Endothelial Dysfunction in Cardiovascular Diseases. Front Biosci (Landmark Ed) 2022, 27: 105

7. Lennicke C, Cochemé HM. Redox metabolism: ROS as specific molecular regulators of cell signaling and function. Mol Cell 2021, 81: 3691–3707

8. Xu S, Ilyas I, Little PJ, Li H, Kamato D, Zheng X, Luo S, et al. Endothelial Dysfunction in Atherosclerotic Cardiovascular Diseases and Beyond: From Mechanism to Pharmacotherapies. Pharmacol Rev 2021, 73: 924–967

9. Chiu T-H, Ku C-W, Ho T-J, Tsai K-L, Yang Y-D, Ou H-C, Chen H-I. Schisanhenol ameliorates oxLDL-caused endothelial dysfunction by inhibiting LOX-1 signaling. Environ Toxicol 2023, 38: 1589–1596

10. Gimbrone MA, García-Cardeña G. Endothelial Cell Dysfunction and the Pathobiology of Atherosclerosis. Circ Res 2016, 118: 620–636

11. Malekmohammad K, Sewell RDE, Rafieian-Kopaei M. Antioxidants and Atherosclerosis: Mechanistic Aspects. Biomolecules 2019, 9: 301

12. Sarmah D, Datta A, Rana N, Suthar P, Gupta V, Kaur H, Ghosh B, et al. SIRT-1/RHOT-1/PGC-1α loop modulates mitochondrial biogenesis and transfer to offer resilience following endovascular stem cell therapy in ischemic stroke. Free Radic Biol Med 2024, 225: 255–274

13. Olmos Y, Sánchez-Gómez FJ, Wild B, García-Quintans N, Cabezudo S, Lamas S, Monsalve M. Sirt1 Regulation of Antioxidant Genes Is Dependent on the Formation of a FoxO3a/PGC-1α Complex. Antioxid Redox Sign 2013, 19: 1507

14. Carrizzo A, Iside C, Nebbioso A, Carafa V, Damato A, Sciarretta S, Frati G, et al. SIRT1 pharmacological activation rescues vascular dysfunction and prevents thrombosis in MTHFR deficiency. Cell Mol Life Sci 2022, 79: 410

15. Liu B, Ou W-C, Fang L, Tian C-W, Xiong Y. Myocyte Enhancer Factor 2A Plays a Central Role in the Regulatory Networks of Cellular Physiopathology. Aging Dis 2023, 14: 331–349

16. Ahmad AA, Meng-Jiao L, Lin J, Guo-Qing L. Myocyte enhancer factor 2 exerts a pivotal role in larval development in Henosepilachna vigintioctopunctata. J Asia-Pac Entomol 2024, 27: 1–9

17. Zhong R, Miao R, Meng J, Wu R, Zhang Y, Zhu D. Acetoacetate promotes muscle cell proliferation via the miR-133b/SRF axis through the Mek-Erk-MEF2 pathway. Acta Biochim Biophys Sin (Shanghai) 2021, 53: 1009–1016

18. Shen X, Zhao X, He H, Zhao J, Wei Y, Chen Y, Han S, et al. Evolutionary conserved circular MEF2A RNAs regulate myogenic differentiation and skeletal muscle development. PLoS Genet 2023, 19: e1010923

19. Cilenti F, Barbiera G, Caronni N, Iodice D, Montaldo E, Barresi S, Lusito E, et al. A PGE2-MEF2A axis enables context-dependent control of inflammatory gene expression. Immunity 2021, 54: 1665–1682.e14

20. Galardi-Castilla M, Fernandez-Aguado I, Suarez T, Sastre L. Mef2A, a homologue of animal Mef2 transcription factors, regulates cell differentiation in Dictyostelium discoideum. BMC Dev Biol 2013, 13: 12

21. Ji X, Chen Z, Wang Q, Li B, Wei Y, Li Y, Lin J, et al. Sphingolipid metabolism controls mammalian heart regeneration. Cell Metab 2024, 36: 839–856.e8

22. Liu B, Pang L, Ji Y, Fang L, Tian CW, Chen J, Chen C, et al. MEF2A Is the Trigger of Resveratrol Exerting Protection on Vascular Endothelial Cell. Front Cardiovasc Med 2022, 8: 775392

23. Li H, Wang F, Guo X, Jiang Y. Decreased MEF2A Expression Regulated by Its Enhancer Methylation Inhibits Autophagy and May Play an Important Role in the Progression of Alzheimer’s Disease. Front Neurosci 2021, 15: 682247

24. Wang L, Fan C, Topol SE, Topol EJ, Wang Q. Mutation of MEF2A in an Inherited Disorder with Features of Coronary Artery Disease. *Science (New York*, NY) 2003, 302: 1578

25. Xu D-L, Tian H-L, Cai W-L, Zheng J, Gao M, Zhang M-X, Zheng Z-T, et al. Novel 6-bp deletion in MEF2A linked to premature coronary artery disease in a large Chinese family. Mol Med Rep 2016, 14: 649

26. Yu J, Yang Y, Xu Z, Lan C, Chen C, Li C, Chen Z, et al. Long Noncoding RNA Ahit Protects Against Cardiac Hypertrophy Through SUZ12 (Suppressor of Zeste 12 Protein Homolog)-Mediated Downregulation of MEF2A (Myocyte Enhancer Factor 2A). Circ Heart Fail 2020, 13: e006525

27. Liu B, Wang L, Jiang W, Xiong Y, Pang L, Zhong Y, Zhang C, et al. Myocyte enhancer factor 2A delays vascular endothelial cell senescence by activating the PI3K/p-Akt/SIRT1 pathway. Aging (Albany NY) 2019, 11: 3768

28. Lee C, Huang C-H. LASAGNA-Search: an integrated web tool for transcription factor binding site search and visualization. Biotechniques 2013, 54: 141–153

29. Singh CK, Chhabra G, Ndiaye MA, Garcia-Peterson LM, Mack NJ, Ahmad N. The Role of Sirtuins in Antioxidant and Redox Signaling. Antioxid Redox Signal 2018, 28: 643–661

30. Di Minno A, Turnu L, Porro B, Squellerio I, Cavalca V, Tremoli E, Di Minno MND. 8-Hydroxy-2-Deoxyguanosine Levels and Cardiovascular Disease: A Systematic Review and Meta-Analysis of the Literature. Antioxid Redox Signal 2016, 24: 548–555

31. Dubois-Deruy E, Peugnet V, Turkieh A, Pinet F. Oxidative Stress in Cardiovascular Diseases. Antioxidants (Basel) 2020, 9: 864

32. Li X, Zou J, Lin A, Chi J, Hao H, Chen H, Liu Z. Oxidative Stress, Endothelial Dysfunction, and N-Acetylcysteine in Type 2 Diabetes Mellitus. Antioxid Redox Signal 2024, 40: 968–989

33. Liu B, Fang L, Mo P, Chen C, Ji Y, Pang L, Chen H, et al. Apoe-knockout induces strong vascular oxidative stress and significant changes in the gene expression profile related to the pathways implicated in redox, inflammation, and endothelial function. Cell Signal 2023

34. Forman HJ, Zhang H. Targeting oxidative stress in disease: promise and limitations of antioxidant therapy. Nat Rev Drug Discov 2021, 20: 689–709

35. Peng X, Sun B, Tang C, Shi C, Xie X, Wang X, Jiang D, et al. HMOX1-LDHB interaction promotes ferroptosis by inducing mitochondrial dysfunction in foamy macrophages during advanced atherosclerosis. Dev Cell 2024: S1534–5807(24)00733-0

36. Guo X, Kesimer M, Tolun G, Zheng X, Xu Q, Lu J, Sheehan JK, et al. The NAD+-dependent protein deacetylase activity of SIRT1 is regulated by its oligomeric status. Sci Rep 2012, 2: 1–7

37. Ministrini S, Puspitasari YM, Beer G, Liberale L, Montecucco F, Camici GG. Sirtuin 1 in Endothelial Dysfunction and Cardiovascular Aging. Front Physiol 2021, 12: 733696

38. Xiao W, Wang R-S, Handy DE, Loscalzo J. NAD(H) and NADP(H) Redox Couples and Cellular Energy Metabolism. Antioxid Redox Signal 2018, 28: 251–272

39. Shelbayeh OA, Arroum T, Morris S, Busch KB. PGC-1α Is a Master Regulator of Mitochondrial Lifecycle and ROS Stress Response. Antioxidants 2023, 12: 1075

40. Chuang YC, Chen SD, Jou SB, Lin TK, Chen SF, Chen NC, Hsu CY. Sirtuin 1 Regulates Mitochondrial Biogenesis and Provides an Endogenous Neuroprotective Mechanism Against Seizure-Induced Neuronal Cell Death in the Hippocampus Following Status Epilepticus. Int J Mol Sci 2019, 20: 3588

41. Batty M, Bennett MR, Yu E. The Role of Oxidative Stress in Atherosclerosis. Cells 2022, 11: 3843

42. Münzel T, Daiber A. Vascular Redox Signaling, Endothelial Nitric Oxide Synthase Uncoupling, and Endothelial Dysfunction in the Setting of Transportation Noise Exposure or Chronic Treatment with Organic Nitrates. Antioxid Redox Signal 2023, 38: 1001–1021

43. Varghese MV, James J, Rafikova O, Rafikov R. Glucose-6-phosphate dehydrogenase deficiency contributes to metabolic abnormality and pulmonary hypertension. Am J Physiol Lung Cell Mol Physiol 2021, 320: L508–L521

44. Bu X, Wu D, Lu X, Yang L, Xu X, Wang J, Tang J. Role of SIRT1/PGC-1α in mitochondrial oxidative stress in autistic spectrum disorder. Neuropsychiatr Dis Treat 2017, 13: 1633–1645

45. Liu J, Jiang J, Qiu J, Wang L, Zhuo J, Wang B, Sun D, et al. Urolithin A protects dopaminergic neurons in experimental models of Parkinson’s disease by promoting mitochondrial biogenesis through the SIRT1/PGC-1α signaling pathway. Food Funct 2022, 13: 375–385

46. Cantó C, Menzies KJ, Auwerx J. NAD(+) Metabolism and the Control of Energy Homeostasis: A Balancing Act between Mitochondria and the Nucleus. Cell Metab 2015, 22: 31–53

47. Imai S, Guarente L. NAD+ and sirtuins in aging and disease. Trends Cell Biol 2014, 24: 464–471

48. Li C, Sun X-N, Chen B-Y, Zeng M-R, Du L-J, Liu T, Gu H-H, et al. Nuclear receptor corepressor 1 represses cardiac hypertrophy. EMBO Mol Med 2019, 11: e9127

49. Ikeda S, He A, Kong SW, Lu J, Bejar R, Bodyak N, Lee K-H, et al. MicroRNA-1 negatively regulates expression of the hypertrophy-associated calmodulin and Mef2a genes. Mol Cell Biol 2009, 29: 2193–2204

50. Shen Y, Zhang T, Jia X, Xi F, Jing W, Wang Y, Huang M, et al. MEF2A, a gene associated with mitochondrial biogenesis, promotes drug resistance in gastric cancer. Biochim Biophys Acta Mol Basis Dis 2025, 1871: 167497

51. Potthoff MJ, Olson EN. MEF2: a central regulator of diverse developmental programs. Development 2007, 134: 4131–4140

52. Xiong Y, Wang L, Jiang W, Pang L, Liu W, Li A, Zhong Y, et al. MEF2A alters the proliferation, inflammation-related gene expression profiles and its silencing induces cellular senescence in human coronary endothelial cells. BMC Mol Biol 2019, 20: 8

53. Li C, Fang F, Wang E, Yang H, Yang X, Wang Q, Si L, et al. Engineering extracellular vesicles derived from endothelial cells sheared by laminar flow for anti-atherosclerotic therapy through reprogramming macrophage. Biomaterials 2025, 314: 122832

54. Lu YW, Martino N, Gerlach BD, Lamar JM, Vincent PA, Adam AP, Schwarz JJ. MEF2 (Myocyte Enhancer Factor 2) Is Essential for Endothelial Homeostasis and the Atheroprotective Gene Expression Program. Arterioscler Thromb Vasc Biol 2021, 41: 1105–1123

55. von Scheidt M, Zhao Y, Kurt Z, Pan C, Zeng L, Yang X, Schunkert H, et al. Applications and Limitations of Mouse Models for Understanding Human Atherosclerosis. Cell Metab 2017, 25: 248–261

56. Xie B, Lin J, Chen X, Zhou X, Zhang Y, Fan M, Xiang J, et al. CircXRN2 suppresses tumor progression driven by histone lactylation through activating the Hippo pathway in human bladder cancer. Mol Cancer 2023, 22: 151

